# Establishment and application of a vesicle extraction method for clinical strains of *Pseudomonas aeruginosa*

**DOI:** 10.1101/2025.11.17.688223

**Authors:** Tania Henriquez, Francesco Santoro, Donata Medaglini, Lucia Pallecchi, Eugenio Paccagnini, Mariangela Gentile, Pietro Lupetti, Chiara Falciani

## Abstract

*Pseudomonas aeruginosa* is a versatile pathogen capable of causing illnesses that range from mild infections to life-threatening conditions. Its virulence is driven by a wide array of factors, among which extracellular vesicles (EVs) have gained recognition as important contributors to its pathogenicity. Despite this, the full scope of their roles remains unclear. A major barrier to EV characterization is the difficulty of vesicle isolation—procedures are often lengthy, yield is low, and specialized equipment is required.

In this study, we assessed the effectiveness of a rapid vesicle extraction method from clinical strains of *P. aeruginosa*. To that end, we first selected and characterized six phenotypically diverse clinical strains of *P. aeruginosa* (two reference strains and 4 clinical isolates, including one strain from a cystic fibrosis patient) and used them to evaluate the vesicle extraction method.

The results obtained through SDS-PAGE analysis, western blot, protein quantification, and TEM indicated the presence of vesicles in all samples; however, it was also possible to observe a large number of contaminants in some of them (mainly LS07 and Z37). Subsequent treatment with enzymes (DNase and/or alginate lyase) allowed for the elimination of the contaminants as observed by electron microscopy.

Our results suggest that the method is suited for the vesicle extraction of clinical isolates of *P. aeruginosa*. The phenotypic complexity of these strains presents challenges that current rapid purification methods are ill-equipped to handle, highlighting the need for improved or alternative approaches.

## 1 Introduction

*Pseudomonas aeruginosa* is a Gram-negative rod-shaped bacterium that can act as an opportunistic pathogen in humans and animals(1–3). This microorganism causes a broad spectrum of diseases, ranging from relatively benign superficial infections—such as those affecting the skin or external ear— to severe and potentially fatal systemic conditions, including pneumonia, urinary tract infections, bacteremia, and sepsis (1–4). Its pathogenicity is particularly pronounced in immunocompromised individuals, such as patients with cystic fibrosis, burn wounds, or those undergoing invasive medical procedures or immunosuppressive therapy (1–4).

The virulence of *P. aeruginosa* is driven by different factors like enzymes, efflux pumps, extracellular vesicles (EVs), among others. EVs, produced from the bacterial membrane(s), play a critical role in spreading virulence factors (5, 6). These vesicles are enriched with a diverse array of bioactive molecules, including proteins, lipids, DNA, RNA, and virulence factors such as exotoxins and enzymes. EVs play different roles in the pathophysiology of *P. aeruginosa* infections by mediating intercellular communication, facilitating horizontal gene transfer, modulating host immune responses, and enhancing bacterial survival and colonization in hostile environments. The unique properties of *P. aeruginosa*-derived EVs have stimulated interest in their potential biotechnological and therapeutic applications (5, 6). Despite these promising developments, several technical challenges hinder the broader application of EVs in both research and clinical contexts. First, their isolation and purification is difficult because of low yields, contamination with non-vesicular material, and high production costs (7). These limitations underscore the need for the development of standardized, scalable, and cost-effective protocols for EV production and characterization.

In addition, it has been observed that clinical strains of *P. aeruginosa*, which are characterized by high phenotypic diversity (8–11), also possess vesicles with diverse protein content and banding patterns on SDS-PAGE gels (12), suggesting that some isolates may be more active than others or more effective as a biotechnological tool. In this context, there is a need to better characterize the wide variety of EVs from clinical strains. However, to the best of our knowledge, there are currently no extraction methods that can be easily applied in a routine clinical laboratory (without an ultracentrifuge), adapted to the clinical phenotypes of *P. aeruginosa*, and that allow for the easy use of vesicles for screening studies.

In recent years, several rapid methodologies, including commercially available extraction kits, have been developed to streamline and simplify laboratory workflows. These methods are designed to enhance efficiency by reducing hands-on time and improving the overall purity of the extracted material. Despite these advantages, the applicability of such rapid protocols can be limited when working with clinical isolates of *P. aeruginosa* (13).

Therefore, here, we assessed the effectiveness of a rapid vesicle extraction method from clinical strains of *P. aeruginosa*. We also analyzed the utility of some treatments to improve the quality of the crude extract of our vesicle samples.

## 2 Material and Methods

### 2.1 Bacterial strains and culture media

*P. aeruginosa* strains were grown in King’s B (KB) medium and maintained on cetrimide agar plates (#70887, Millipore, prepared according to the manufacturer’s instructions). For long-term storage, they were kept in glycerol stocks at −80°C. A complete list of the strains used in this study can be found in Table 1. All strains were cultured at 37°C, unless otherwise specified. King B medium (prepared as previously described (14)) was used to grow the strains for vesicle isolation.

**Table 1.** List of strains used in this study.

| Name | Origin (sample type) | Reference |
| --- | --- | --- |
| PAO1 | Wound | (15) |
| ATCC27853 | Blood | (16) |
| LS03 | Sputum | (13) |
| LS06 | Urine | (13) |
| LS07 | Cerebrospinal fluid | (13) |
| Z37 | Sputum (cystic fibrosis patient) | (17) |

### 2.2 Vesicle isolation

Strains were grown in 2.5 ml of KB medium at 37°C and used to inoculate 500 ml of KB medium. The cultures were incubated overnight at 37°C with continuous shaking until reaching an OD600 of 0.790-1.21. Later, the cultures were centrifuged twice at 3214 x *g* for 30 min at 4°C, and supernatants were passed through 0.45 and 0.22 μm filters. TFF cassettes (Vivaflow SU, 100 kDa, Sartorius, VF-S050H0100-IV) were used to concentrate the samples. Samples were recovered with the addition of 30 ml of 10 mM HEPES 0.85% (w/v) NaCl (final volume of 50 ml). Later, the samples were centrifuged using a Vivaspin 100 kDa device and recovered in 10 mM HEPES 0.85% (generating a final volume of 1000-1500 μl). A final centrifugation was performed to remove flagellar proteins at 16 000 x g for 30 min (18). The samples were aliquoted and stored at −20°C.

### 2.3 Phenotypic characterization of strains

For pigment production (including pyoverdine) and mucoid phenotype detection, strains were streaked on cetrimide and King B agar plates, and incubated at RT for 13 days.

### 2.4 DNA gel electrophoresis

A 2% agarose gel was prepared in 1X TBE buffer plus 1X GelStar (Lonza, # 50535). Vesicle samples were loaded on the gel (using GelPilot loading dye 5X, QIAGEN, # 239901) and an electrophoresis was performed at RT for 30 min at 85 V. Finally, a picture of the gel was taken using a 50 MP camera (moto g31(w), XT2173-3).

### 2.5 SDS-PAGE and western blot experiments

Vesicle samples were used to run two identical SDS-PAGE gels in parallel (12% mPAGE precast gels (#MP12W15, Merck) in 1× MES SDS, 100 V for 60 min). One of the gels was stained with Thermo Scientific Imperial Protein Stain (#24615, Thermo Scientific) for two hours and destained overnight with water. The second gel was transferred to a nitrocellulose membrane using a Trans-Blot Turbo Mini 0.2 μm Nitrocellulose Transfer Pack (#1704158, Bio Rad), according to the manufacturer’s instructions. The membrane was then blocked (3% BSA in TBS) and incubated on a rocker shaker for 1 h at RT. After that, primary antibody solution (containing anti-OprF, Thermo Fisher #PA5-117553, diluted 1:2,500 in TBS +0.05% Tween, 3% BSA) was added and incubated 1 h at RT on a rocker shaker. Later, the solution was removed, and the membrane was washed three times with TBS + 0.05% Tween (10 min each time) on a rocker shaker at RT. A secondary antibody solution (containing goat Anti-Rabbit IgG antibody, Alkaline Phosphatase conjugate, Sigma-Aldrich # AP132A diluted 1:10,000 in TBS +5% skimmed milk) was added and incubated for 1 h at RT on a rocker shaker. Again, the membrane was washed three times with TBS + 0.05% Tween (10 min each time) on a rocker shaker at RT. Finally, the buffer was discarded, and BCIP®/NBT (Sigma-Aldrich # B1911) was added on top of the membrane (according to the manufacturer’s instructions), incubated for 10 min at room temperature and photographed.

### 2.6 Transmission electron microscopy

For electron microscopy analysis, 3 μL of each sample was loaded onto formvar-coated 300 mesh Cu grids (Ted Pella Inc., Redding, CA) for 2 min at RT. After removing the excess of sample with filter paper, 3 μL of 1% uranyl acetate (Polysciences Inc., Warrington, PA) in distilled water was added for 30 s, blotted again with filter paper and air dried. Finally, the samples were analyzed using a Thermo Fisher Scientific Tecnai G2 Spirit 120 kV transmission electron microscope (equipped with a TVIPS TemCam-F216 CMOS camera).

### 2.7 Protein quantification

For protein quantification, bicinchoninic Protein Assay Kit for dilute samples (Euroclone, # EMP015480) was used according to manufacturer’s instructions. Briefly, working reagent was prepared by mixing 25 parts of reagent A, 24 parts of Reagent B and 1 part of Reagent C. In parallel, a standard curve using BSA as sample was prepared. Later, 150 μl of working reagent was mixed with 150 μl of sample, standard or blank, and incubated at 37°C for 2 h. Then, the samples were cooled down at RT, absorbance was measured at 562 nm and a plot was generated using the reading of each standard vs. its concentration (after correcting for the value of the blank). Finally, the protein concentration of each unknown sample was calculated using the calibration plot.

### 2.8 Enzymatic digestion

Samples were treated, when needed, with DNase (#EN0521, Thermo Scientific) and/or Alginate lyase (#A1603, Sigma-Aldrich). Briefly, for DNase I treatment, 26 μl vesicle sample were digested with 1 U DNase in presence of 1X reaction buffer with MgCl_2_. The samples were incubated at 37 °C for 30 min and the reaction was stopped at 65 °C for 10 min after the addition of 1 μl 50 mM EDTA. Later, the sample was centrifuged using an Amicon Ultra 0.5 ml (100 kDa) filter and recovered with 150 μl 10 mM HEPES 0.85 NaCl. For alginate lyase, the samples were treated according to manufacturer instructions with a small modification (12 units of enzyme were added to the reaction). Incubation was performed at 37°C for 10 min. Reaction was terminated by addition of 0.1 N NaOH and the samples were then concentrated with Amicon Ultra 0.5 ml (100 kDa) filter and recover with 150 μl 10 mM HEPES 0.85 NaCl.

## 3 Results and discussion

### 3.1 Strain characterization

We selected six clinical isolates of *P. aeruginosa* (including one strain from a patient with cystic fibrosis, Table 1) from a group of strains that we previously characterized (13, 17). We then analyzed them for pigment production and mucoid phenotype in cetrimide agar. To that end, the strains were grown 13 days at RT on cetrimide agar plates, a selective medium for *Pseudomonas* that induces the production of pigments such as pyoverdine and pyocyanin (19). Our results indicated that after long-term incubation, the strains produced different pigments (Figure 1) and only Z37 strain showed a mucoid phenotype in this agar (Figure 1). This finding reinforces our previous characterization and supports the fact that these strains are phenotypically diverse, and representative of the variability described for clinical strains of *P. aeruginosa* (8, 13).

**Figure 1.**
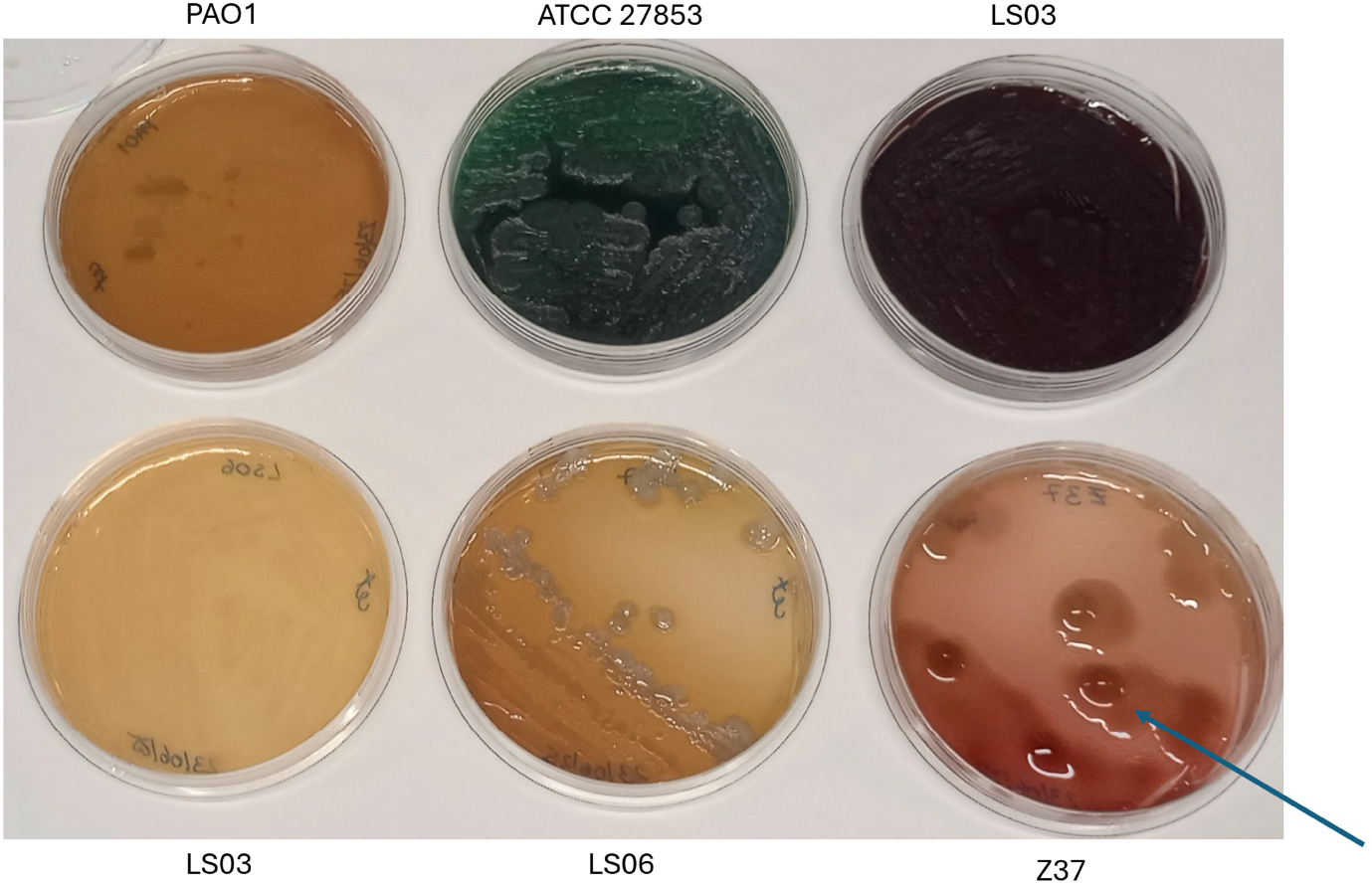
Mucoid phenotype and pigment production analysis. *P. aeruginosa* strains were grown on cetrimide agar and monitored for 13 days at room temperature and photographed. Blue arrow indicates mucoid colony of Z37 strain.

### 3.2 Vesicle extraction and characterization

Previously, we tested a commercial kit for the isolation of vesicles from *P. aeruginosa* clinical strains with little success (13). Based on our previous experience, we decided to design a protocol for the extraction of vesicles from clinical isolates. To that end, we adapted/modified protocols reported by other groups for *Pseudomonas* or for other microorganisms (18, 20–22). The rationale for the protocol was to use equipment that is normally present in microbiology laboratories and/or that can be easily acquired (Figure 2). In this context, small volumes of culture were preferred as they are much more easily centrifuged in comparison to liters. Briefly, an inoculum was grown during the day and used to inoculate a flask containing 500 ml of KB, which was then incubated overnight. The sample was centrifuged and filtered several times (using 0.45 and 0.22 μm filters). At the end of the ultrafiltration, a vesicle crude extract was obtained. Finally, the sample was centrifuged at 16000 x g for 30 minutes in order to remove flagellar proteins as suggested by previous works (18).

**Figure 2.**
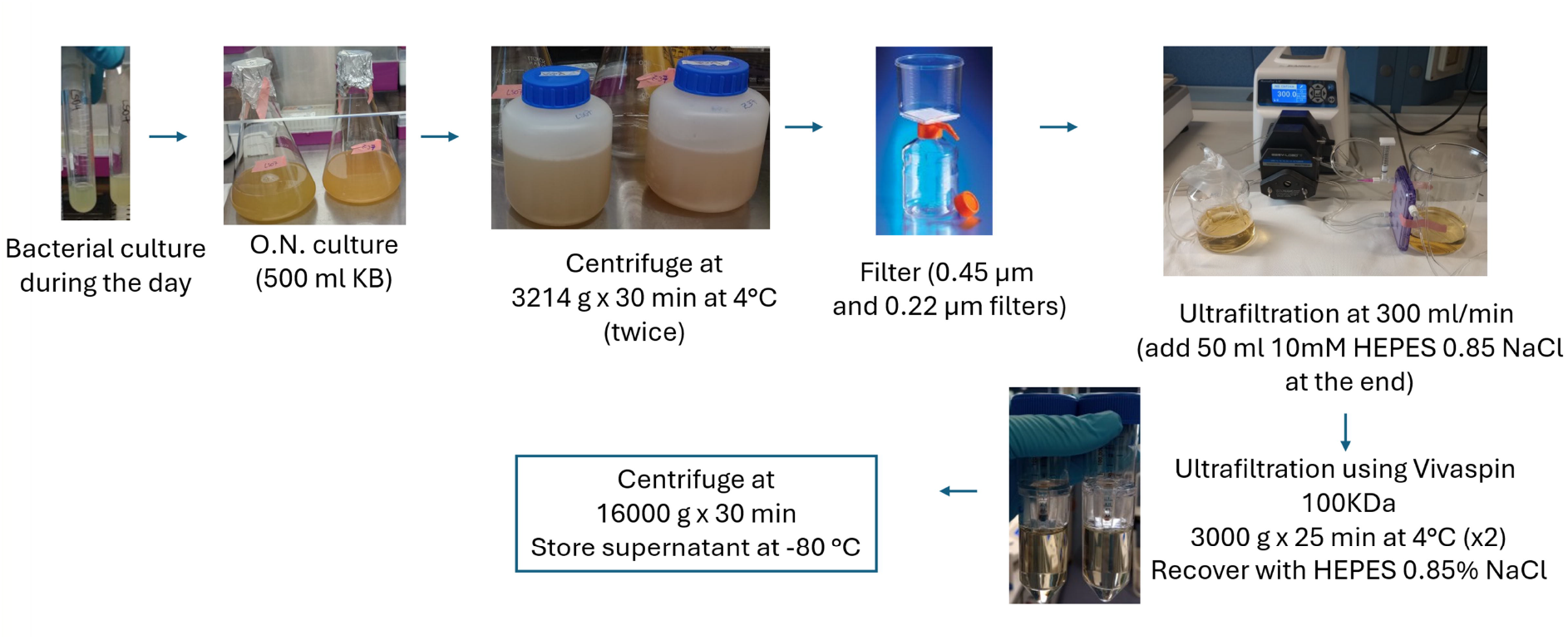
Vesicle extraction method.

A pilot experiment was conducted with *P. aeruginosa* PAO1 strain. Vesicle sample was analyzed by SDS-PAGE and WB using an antibody against OprF porin, an important membrane protein of *P. aeruginosa* (23, 24) (Figure 3A). SDS-PAGE showed several bands for both samples (PAO1 sample before and after final centrifugation to remove contaminants) and WB showed positive signal for OprF porin, which is compatible with the presence of vesicles in the samples.

**Figure 3.**
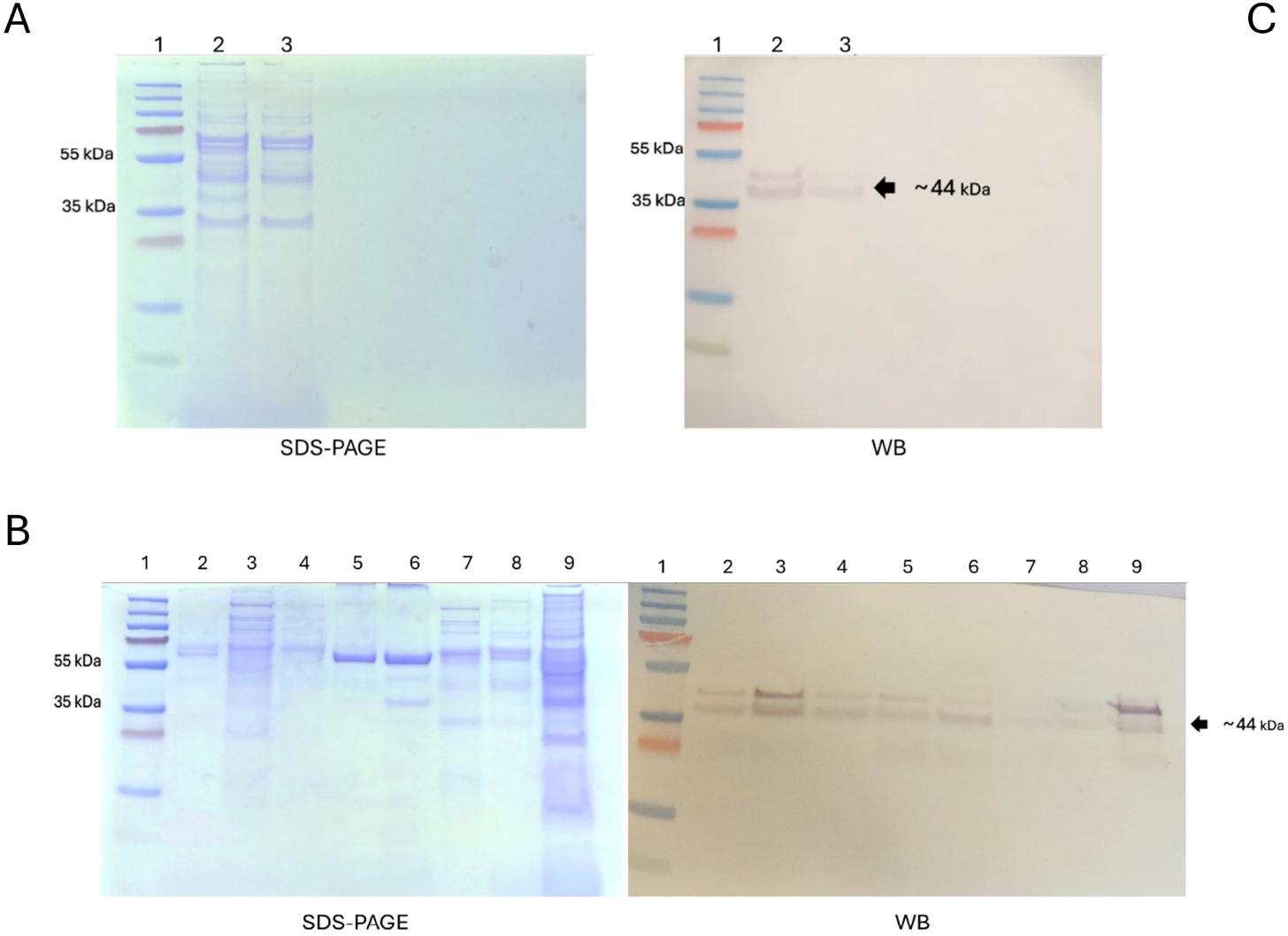
Vesicle analysis. (A) vesicles from PAO1 strain were analyzed by SDS-PAGE (left) and western blot, WB (right). For SDS-PAGE, Imperial Protein stain was used. For WB, an antibody against OprF porin conjugated with alkaline phosphatase was used. For both images, 1: PageRuler™ Prestained Protein Ladder; 2: PAO1 vesicles – crude extract; 3: PAO1 vesicles after last centrifugation to remove flagella. (B) Vesicle samples from clinical strains were analyzed by SDS-PAGE (left) and WB (right). For SDS-PAGE Imperial Protein was used and for WB, antibody against OprF porin. For both images, 1: PageRuler™ Prestained Protein Ladder; 2: PAO1; 3: ATCC 27852; 4: LS03; 5:LS06; 6:LS07; 7: Z37; 8: PAO1 (final sample – pilot experiment); 9: PAO1 pellet (pilot experiment).

Then, the same protocol was used to extract vesicles from all six *P. aeruginosa* strains. Samples were analyzed by SDS-PAGE, WB (Figure 3B) and TEM (Figure 4). Protein quantification indicated that all samples had a concentration above 400 μg/ml (between 432 and 1726 μg/ml, Table 2). Our results showed that all the vesicle samples had a different protein pattern in SDS-PAGE, which is in agreement with previous reports (12). Also, all the samples were positive for OprF, although Z37 had a signal level much lower than the others. In parallel, TEM analysis showed vesicles in samples PAO1, ATCC 27853, LS03 and LS06 (Figure 4). However, the images also showed the presence of flagellar proteins, indicating that the final centrifugation step reduced these contaminants but was not enough to eliminate them. Additionally, it was not possible to clearly observe vesicles in samples LS07 and Z37, as they contained some contaminants (Figure 4). These contaminants could be the reason for the low signal detected for OprF by WB.

**Table 2.** Protein concentration of different vesicle samples.

| Sample | Concentration (µg/ml) |
| --- | --- |
| PAO1 | 567.3504 |
| ATCC 27853 | 1092.137 |
| LS03 | 432.3077 |
| LS06 | 485.2991 |
| LS07 | 1726.325 |
| Z37 | 693.8462 |

**Figure 4.**
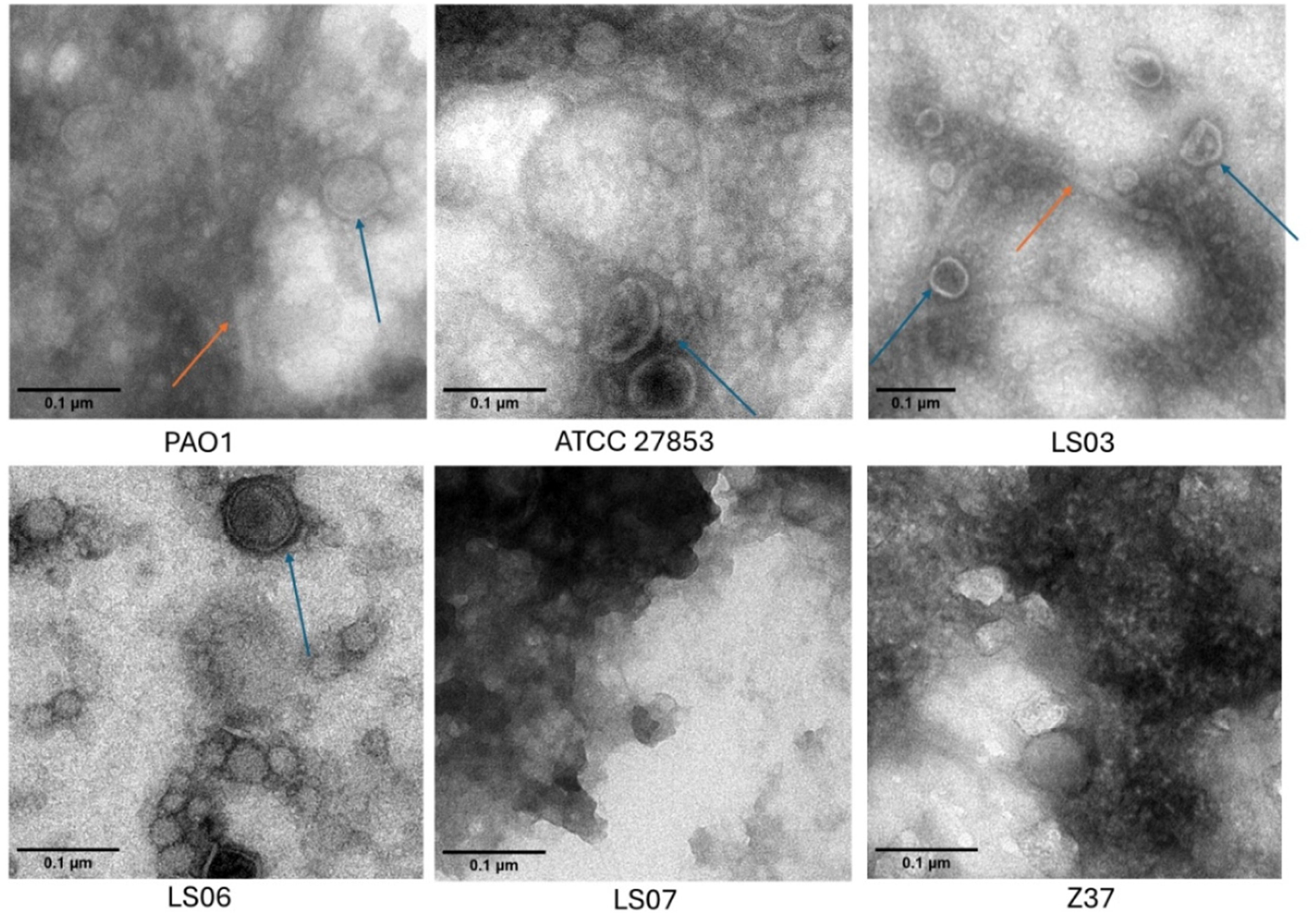
Electron microscopy images of vesicle samples. Blue arrows indicate some extracellular vesicles and orange arrows indicate flagellar proteins. Samples LS07 and Z37 showed a high number of unidentified contaminants.

In this context, we treated samples LS07 and Z37 with alginate lyase, as alginate has been described as an important component of the *Pseudomonas* capsular exopolysaccharide and a driver of the mucoid phenotype (25).

Our TEM results indicated that alginate lyase treatment was successful in removing contaminants from sample LS07 (Figure 5A), however sample Z37 still contained additional particles not related to vesicles (data not shown). Since agarose gel electrophoresis revealed the presence of genomic DNA and what could be plasmid DNA (Figure S1), it was decided to additionally treat sample Z37 with DNase I. Our results showed that the combined enzymatic treatment was useful to obtain a cleaner Z37 sample according to TEM (Figure 5B). However, it is important to mention that several vesicles seemed to be damaged or aggregated, suggesting that this longer protocol could be stressful for them. In addition, the individual vesicles observed were, on average, smaller than those of the other strains.

**Figure 5.**
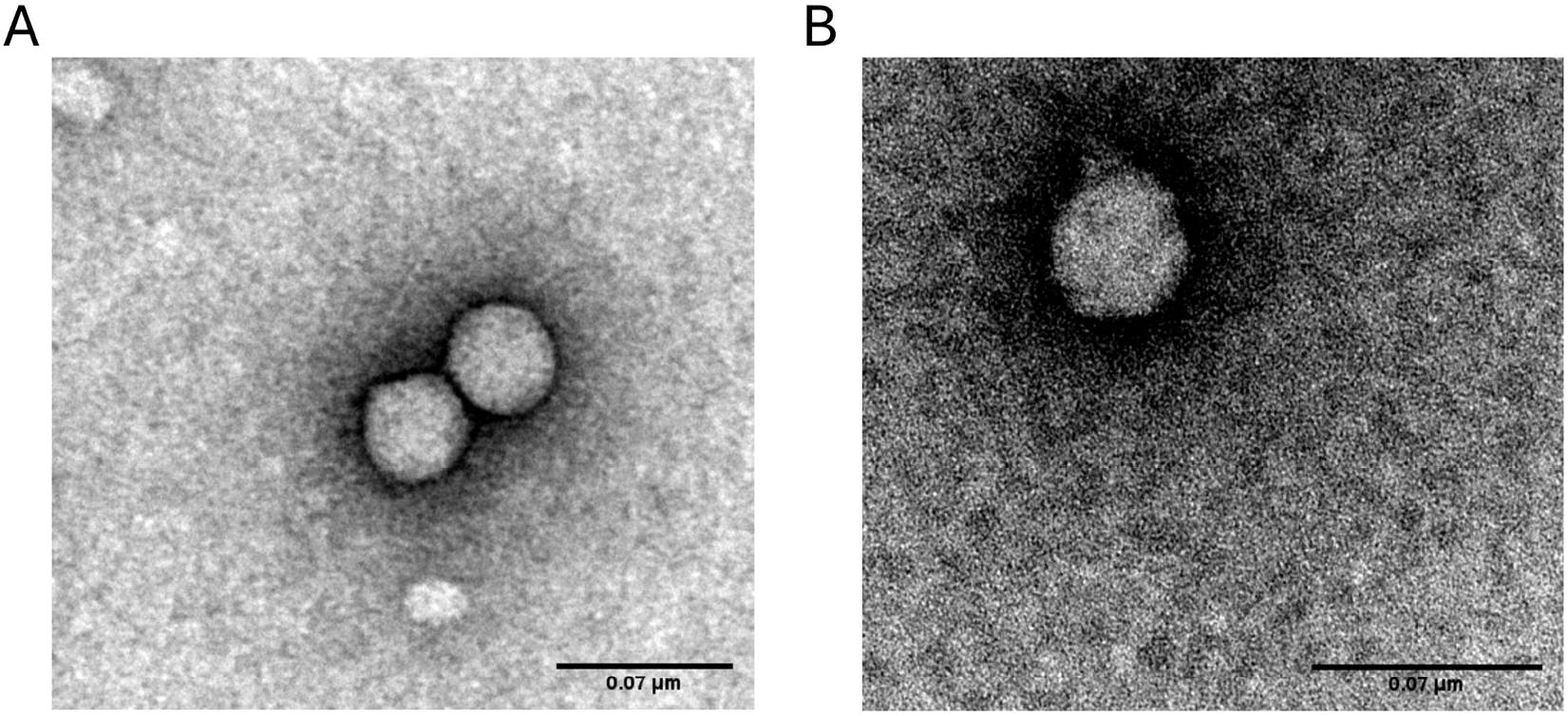
Electron microscopy images of alginate lyase treated vesicle samples. Samples (A) LS07 and (B) Z37 were treated with 12 units of alginate lyase for 10 min (and 1 U DNase for 30 min in the case of 37) and analyzed by TEM.

Nevertheless, the results indicated that it was possible to clean the samples of most of the interferents observed in the TEM images, suggesting the usefulness of the protocol and the possibility of using these samples for further functional studies.

## 4 Conclusion

In summary, the phenotypic complexity of these strains presents challenges that conventional purification methods are unable to address. In this context, it is expected that the method designed in this study will aid in the rapid and efficient extraction of EVs in clinical laboratories, allowing these samples to be used for functional screening. In this way, only preparations with relevant activities could then be purified using conventional protocols (such as density gradient ultracentrifugation), if required.

In conclusion, this protocol is expected to support further investigations into how distinct vesicle populations derived from clinical *P. aeruginosa* strains contribute to bacterial physiology and pathogenicity.

## 7 Acknowledgments

We thank Clelia Cortese for her help with some experiments.

## 9 Supplemental material

Figure S1. Agarose electrophoresis of vesicle samples

