## Supplemental material for "Establishment and application of a vesicle extraction method for clinical strains of *Pseudomonas aeruginosa*"

Methods and Protocols article

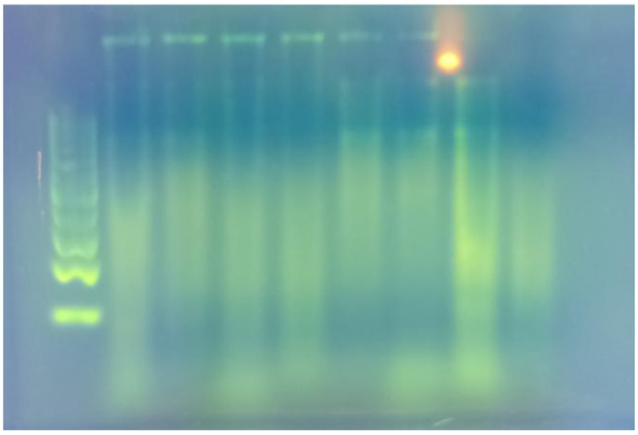


**Figure S1.** Agarose electrophoresis of vesicle samples. Extracellular vesicles were used as a sample for a 2% agarose gel electrophoresis with 1X GelStar dye. 1: DNA marker; 2: PAO1 (pilot experiment); 3: PAO1; 4: ATCC 27853; 5: LS03; 6: LS6; 7: LS07; 8: Z37; 9: Z37 after DNase treatment.
